# Adaptive Gain model for predicting auditory brain activity in mice outperforms standard methods for predicting cortical speech tracking in human EEG

**DOI:** 10.64898/2026.02.12.705421

**Authors:** Adèle Simon, Adrian N. L. Sahani, Maria Chait, Jennifer F. Linden

## Abstract

Human brain activity tracks slow fluctuations in continuous sound, including speech, and modelling this relationship provides insight into how the brain encodes naturalistic auditory input. Traditional approaches to modelling brain tracking of speech with linear regression often use the amplitude envelope of the stimulus as a regressor, implicitly assuming that neural responses scale linearly with sound intensity. However, previous studies of central auditory processing in both animals and humans have shown that adaptive mechanisms adjust auditory responses based on recent sound history. Here, we find that a simple nonlinear transformation of the stimulus envelope, derived from studies of mouse auditory thalamic responses to temporally varying sounds, significantly improves modelling of human cortical speech tracking. This Adaptive Gain transformation essentially normalizes sound level by its recent context. Improvements in modelling of cortical speech tracking using the Adaptive Gain stimulus representation are robust across experiments, participant groups, and languages, indicating that short-timescale adaptation to recent sound level is a general feature of auditory processing. We further show that the optimal adaptation time constant for human cortical responses to continuous speech is approximately 50–100 ms, longer than previously observed for auditory thalamic responses to temporally varying sounds in mice. In summary, the Adaptive Gain transformation is a mathematically simple alternative to the standard envelope representation that captures dynamic adaptation in auditory processing and reliably improves prediction of human cortical EEG signals during listening to continuous speech.

**Significance Statement:** Standard models of human brain responses to continuous speech use the raw stimulus envelope as a predictor of EEG responses, implicitly assuming stationary sensitivity to sound intensity. Here, we use instead a simple nonlinear transformation of the stimulus envelope, Adaptive Gain, which normalizes sound level based on recent sound history over tens to hundreds of milliseconds. Originally derived from studies of auditory processing in mice, the Adaptive Gain transformation robustly improves prediction of EEG responses to continuous speech in humans across experiments, participant groups, and languages. Results demonstrate that dynamic adaptation to sound level is a core feature of auditory processing, and can be approximated in a mathematically simple form to improve prediction of cortical speech tracking.

## 1 Introduction

In humans listening to continuous speech, cortical EEG activity tracks slow fluctuations in the auditory signal. This “cortical speech tracking” phenomenon is believed to reflect the aggregation of successive evoked responses to transient acoustic events within the ongoing speech stream (Aiken and Picton, 2008; Oganian et al., 2023). The amplitude envelope representation of speech captures power variations that can drive such evoked responses, and several studies have highlighted a link between the envelope of an incoming sound and the neural response measured using EEG (Etard and Reichenbach, 2019; Simon et al., 2024; Decruy et al., 2019). This link can be analyzed by fitting linear “Temporal Response Function” (TRF) models relating the audio representation to the neural response (Lalor et al., 2009; Crosse et al., 2016) and assessing how accurately such models predict neural responses to novel audio inputs (Brodbeck and Simon, 2020; Meyer et al., 2017). Because TRFs are linear models, they cannot directly capture nonlinear mechanisms, but the impact of nonlinear processes on cortical speech tracking can be studied by embedding those nonlinearities in the stimulus representation and comparing how well the resulting models predict neural responses (Drennan and Lalor, 2019; Lindboom et al., 2023; Biesmans et al., 2016; Williamson et al., 2016; Kulasingham et al., 2024; Shan et al., 2024; Maddox and Lee, 2018).

The most commonly used regressor for TRF modelling is the amplitude envelope, based on substantial evidence for the central role of the envelope in speech perception (Shannon et al., 1995; Aiken and Picton, 2008; Smith et al., 2002; Moon et al., 2014). However, using the envelope as a stimulus representation means assuming that the cortical response depends solely on the absolute value of speech amplitude at a given moment. This assumption may be incorrect, as it overlooks adaptive mechanisms that adjust neural responses to transient events based on their context (Lohse et al., 2020; Willmore and King, 2023; Ahrens et al., 2008). Such adaptive mechanisms have been documented across multiple stages of the auditory pathway (Shamma and Fritz, 2014; Pérez-González and Malmierca, 2014), including at both subcortical levels (Willmore and King, 2023; Kvale and Schreiner, 2004; Dean et al., 2005) and cortical levels (Williamson et al., 2016; Akritas et al., 2024; Ahrens et al., 2008; Shechter and Depireux, 2006; Ulanovsky et al., 2004; Werner-Reiss et al., 2006; David and Shamma, 2013), and evidence for adaptive mechanisms has also been noted at the perceptual level (Simpson et al., 2014; Khalighinejad et al., 2019). Adaptation to auditory context is thought to be critical for sound perception in noisy environments (King and Walker, 2020; Auerbach and Gritton, 2022), including speech-in-noise situations (Ding and Simon, 2013).

Here, we asked whether a simple adaptive nonlinear function of the stimulus envelope previously derived from a model of temporal processing in the auditory thalamus of anesthetized mice (Anderson and Linden, 2016) could be used to improve cortical tracking of continuous speech in awake humans.

The “Adaptive Gain” transformation essentially normalizes the amplitude envelope by recent sound level weighted by a function that decays exponentially in preceding time. Adaptive Gain can be interpreted as representing the dynamic responsiveness of the central auditory system to sound level. We find that even when applied only to the broadband stimulus envelope and with parameters optimized originally for modelling neural population responses in the auditory thalamus of anesthetized mice, the Adaptive Gain representation improves human cortical speech tracking across experiments, participant groups, and languages. We also identify optimized Adaptive Gain parameters for human studies that enhance cortical speech tracking even further.

The results demonstrate that cortical speech tracking is strongly influenced by nonlinear adaptation to sound level on a timescale of tens to hundreds of milliseconds, and that a simple, easily applied nonlinear transformation of the stimulus envelope enables improved prediction of human EEG responses to continuous speech.

## 2 Methods

### 2.1 Data

#### 2.1.1 Dataset description

Two separate datasets (Simon et al., 2022; Etard and Reichenbach, 2022) were used for the present study. Both were publicly available datasets. They were chosen based on the following criteria: (i) EEG was recorded during speech listening, with continuous segments lasting at least 30 seconds; (ii) recording was done during listening to clean speech, without additional noise; (iii) participants did not report any hearing or neurological disorders; (iv) the raw EEG data was available; and (v) the original audio used in the experiments was available.

The first dataset anaylsed here (Simon et al., 2022) is a subset of the data collected and used in Simon et al. (2024). It consists of EEG recordings from 21 native Danish speakers listening to audiobooks in Danish and in Finnish without background noise for approximately 6 minutes for each language (divided into 70-second trials). Audiobooks were played on loudspeakers at 60 dBA measured at the listening position. Participants’ cortical responses while listening were recorded continuously at 2400 Hz with a g.HIamp-Research (g.tec Medical Engineering GmbH, Austria) system and active electrodes positioned according to the standard 10-20 system.

The second dataset (Etard and Reichenbach, 2022) is a subset of the collected data used for Etard et al. (2019); Etard and Reichenbach (2019). It comprises data recorded in 18 native English speakers listening to 10 minutes of natural continuous English speech and 10 minutes of natural continuous Dutch speech without background noise, through Etymotic ER-3C insert tube earphones at 76dBA. Participants’ cortical responses were continuously recorded at 1 kHz through a 64-channel scalp EEG system using active electrodes (actiCAP, BrainProducts, Germany) and a multi-channel EEG amplifier (actiCHamp, BrainProducts, Germany). The electrodes were positioned according to the standard 10-20 system and referenced to the right earlobe.

Both datasets included data recorded while participants listened to speech in a language they understood and also to speech in a foreign language they did not understand. We focus here primarily on analysis of data for understood speech, to avoid effects due to language understanding or variation in attention. However, we also analyzed data for not-understood speech and obtained similar results, as shown at the end of the Results section.

#### 2.1.2 EEG preprocessing

All EEG data were preprocessed using custom-written Matlab code (R2022a; The MathWorks, Natick, MA, USA) combined with the EEGLAB toolbox (Delorme and Makeig, 2004). The EEG data were rereferenced to an average reference. The data were filtered with first a 3rd-order high-pass Butterworth filter with a cutoff frequency of 0.5 Hz, then a 5th-order low-pass Butterworth filter with a cutoff frequency of 15 Hz (Crosse et al., 2021). Both filters were implemented using the *filtfilt* function in Matlab, to ensure zero phase distortion (de Cheveigné and Nelken, 2019). Data were downsampled to 200 Hz to reduce processing time and separated into trials. The first and last 5 seconds of each trial were removed to exclude the onsets and offsets of the speech stimuli (Crosse et al., 2021). Finally, EEG signals were z-score-normalized before estimation of Temporal Response Functions relating the EEG to different audio representations.

### 2.2 Audio Representations

Three audio representations were extracted from the original audio stimuli and used as regressors for TRF modelling: Envelope, Logarithmic Envelope (LogEnv), and Adaptive Gain.

After extraction, all audio representations were downsampled to 200 Hz to match the EEG sampling frequency using the *decimate* function in MATLAB (version R2022a), which includes an 8th-order low-pass Chebyshev Type I infinite impulse response anti-aliasing filter with a cut-off frequency of 79 Hz.As the absolute values differ between the three audio representations, all audio representations were z-score-normalized to standardize them relative to overall temporal variability. Examples of the different audio representations of noise and speech are shown in Figure 1.

**Figure 1:**
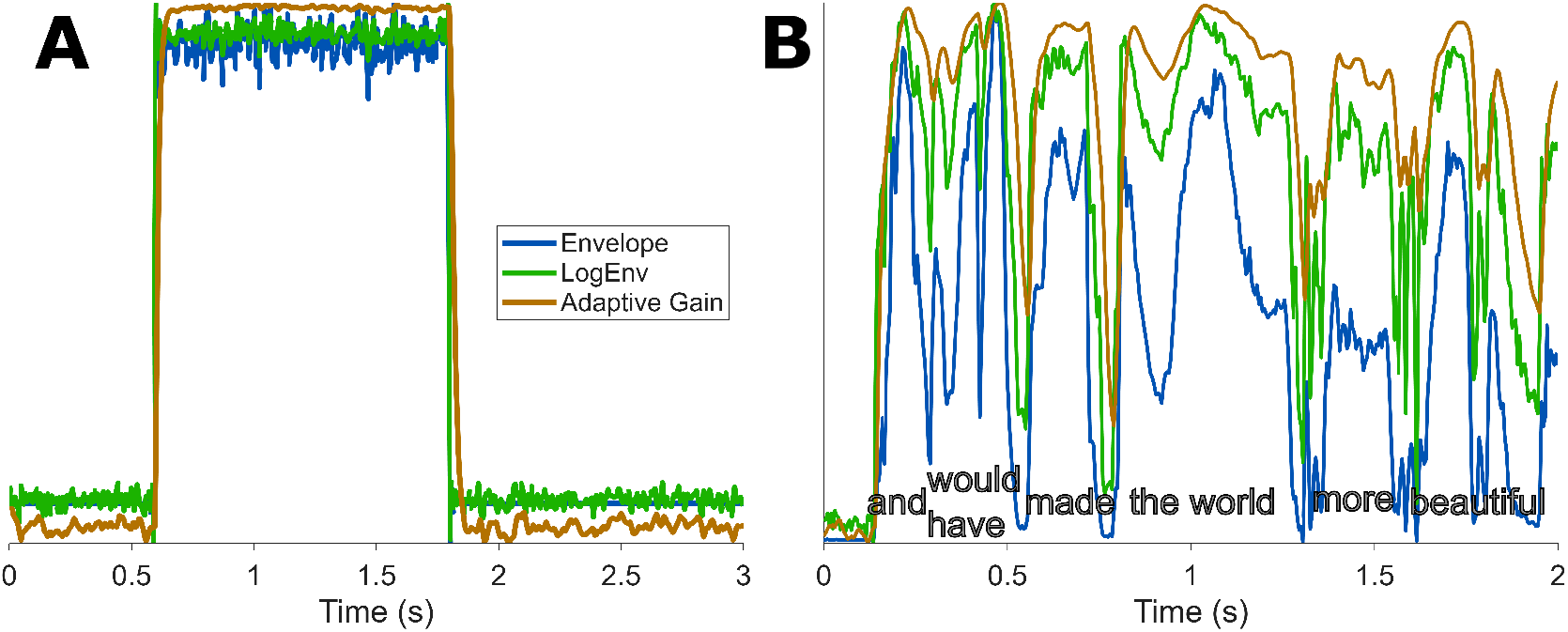
Examples of the audio representations compared in the current study. For visualization purposes, Adaptive Gain values were vertically flipped (i.e., multiplied by −1 and shifted up) and then all three audio representations were scaled to the same range, so that temporal variations in the different representations could be more easily compared. **A**, Envelope, LogEnv and Adaptive Gain representations of a burst of white noise in silence. **B**, Envelope, LogEnv and Adaptive Gain representations of two seconds of continuous speech.

#### 2.1.1 Envelope

The envelope corresponds to the slower energy fluctuations that are tracked by the auditory cortex (Lalor and Foxe, 2010; Ahissar et al., 2001) and are known to be crucial for speech intelligibility (Shannon et al., 1995). It has been shown that cortical speech tracking is dominated by the processing of the amplitude envelope (Prinsloo and Lalor, 2022). The envelope is also the most commonly used audio representation in speech tracking research (Etard and Reichenbach, 2019; Simon et al., 2024; Zuk et al., 2021; Decruy et al., 2019). Therefore, the envelope served as a reference for assessing whether other audio representations could enhance TRF modelling performance.

The envelope was extracted from the speech signals by applying a full-wave rectification to the original sound pressure waveform *s*(*t*) as:

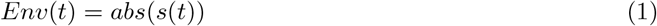

No additional low-pass filtering was applied following the full-wave rectification, as a low-pass filter (with a cut-off frequency of 79 Hz) was applied as a part of the downsampling.

#### 2.2.2 Logarithmic Envelope (LogEnv)

The Adaptive Gain computation relies on the broadband envelope of the audio signal, logarithmically scaled, extracted as follows:

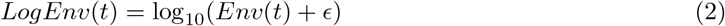

where Env(t) is the envelope of the audio signal as defined in the previous subsection, and *ϵ* is a small positive constant included to ensure that the argument of the logarithm remains positive. In both human and animal studies, it has been shown that central auditory encoding of the intensity of an auditory signal is better described using logarithmic than linear representations of sound intensity (Escabí et al., 2003; Ahrens et al., 2008; Rahman et al., 2020; Aiken and Picton, 2008). However, previous work failed to find an improvement in speech tracking for the logarithmic representation compared to the envelope (Drennan and Lalor, 2019; Biesmans et al., 2016). As the adaptive gain computation incorporates a logarithmic transformation of the envelope, we included the LogEnv representation among the audio representations compared to the Adaptive Gain representation in order to determine the extent to which logarithmic transformation alone might account for any improvements in speech tracking performance.

#### 2.2.3 Adaptive Gain

The Adaptive Gain transformation can interpreted as capturing dynamic responsiveness of the central auditory system to sound level on a millisecond timescale; it is high when sound levels have recently been low, and low when sound levels have recently been high. Computation of Adaptive Gain is a key step in simple 6-parameter “onset-offset” model of auditory temporal processing originally developed to predict neural population activity evoked by gap-in-noise stimuli in the auditory thalamus of anaesthesized mice (Anderson and Linden, 2016). The Adaptive Gain calculation is not intended to model specific physiological or cellular processes but to describe the adaptive intensity gain control performed by the auditory pathway through a nonlinear normalization (Carandini and Heeger, 2012). The Adaptive Gain function *AG*(*t*) can be computed as follows:

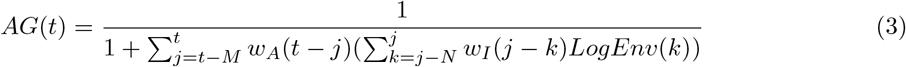

where *LogEnv*(*k*) is the logarithmic envelope transformation of the audio signal as defined in the previous subsection. *w*_*I*_ is the integration window defined by:

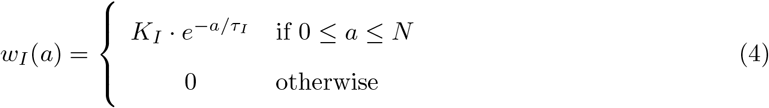

where *N* is the maximum duration of the integration window and is set to 5*τ*_*I*_ to ensure *>* 99% decay of the exponential at the earliest time points falling within the window. *K*_*I*_ is chosen such that 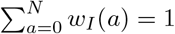.

*w*_*A*_ is the adaptation window defined in a similar manner:

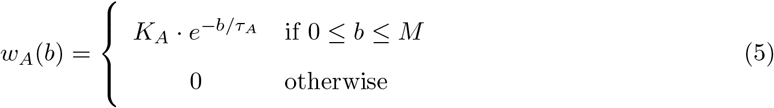

where *M* is the maximum duration of the integration window and is set to 5*τ*_*A*_. *K*_*A*_ is defined to have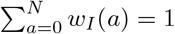.

The two successive convolutions *w*_*I*_ *∗ w*_*A*_ within the Adaptive Gain calculation can be simplified to the following mathematical form (see Supplementary Information for details and further explanation):

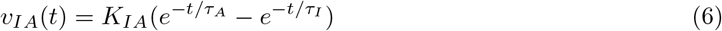

with *τ*_*A*_ and *τ*_*I*_ as defined above, and *K*_*IA*_ a single constant defined such that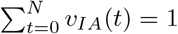. This simplification produces a single smooth weighting function of time preceding the current moment in the sound signal. The *v*_*IA*_(*t*) function is zero at *t* = 0 and rises rapidly to a peak at a preceding time point, then falls more gradually across time steps further into the past.

As can be seen in Equation 6, the integration and adaptation windows *τ*_*I*_ and *τ*_*A*_ are mathematically equivalent and interchangeable in the computation of the Adaptive Gain. They were, however, designed to describe distinct operations in the larger model of temporal processing developed in Anderson and Linden (2016) to predict neural population responses to gap-in-noise stimuli in the mouse auditory thalamus. Therefore, we have retained the original labelling of integration and adaptation time constants in our descriptions here, even though the distinction is irrelevant for the Adaptive Gain computation in isolation.

For our initial analysis of human cortical speech tracking using the Adaptive Gain representation of continuous speech, the values of the variables defining the length of the integration and adaptation windows were fixed to values similar to those that had been identified as optimal for predicting neural population responses to gap-in-noise stimuli in the auditory thalamus of anesthetized mice (Anderson and Linden, 2016) (*τ*_*A mice*_ = 10 ms and *τ*_*I mice*_ = 6 ms). As *τ*_*I*_ was originally defined to be an integration window representing more fundamental limits for temporal integration of the stimulus envelope (e.g., on the order of a few milliseconds, close to the limits of gap-in-noise detection for both mice and humans), we set *τ*_*I human*_ = *τ*_*I mice*_ = 5 ms, rounding the original mouse-optimized value of 6 ms in Anderson and Linden (2016) down to 5 ms to align with the 200 Hz downsampled EEG sample rate used here. The adaptation time constant *τ*_*A*_ is theoretically less constrained, and the mouse-optimized values may not be optimal for modelling adaptation at the cortical level in humans. Thus, a second step in the analysis involved an estimation of the optimal *τ*_*A human*_ values. Optimization of the adaptation parameter was done with a grid search to find the values of *τ*_*A*_ that yielded the best performance for a forward TRF model. The values tested in this grid search were *τ* = 5, 10, 25, 50, 75, 100, 250, and 500 ms. Increasing *τ*_*A*_ will flatten the shape of the *v*_*IA*_ function, which increases the amount of previous data included in the smoothing. Increasing *τ*_*A*_ also introduces a delay in the peak of the *v*_*IA*_ function, as illustrated in Supplementary Information.

The *v*_*IA*_ formulation allows us to rewrite the equation for the Adaptive Gain function *AG*(*t*) more simply in terms of a convolution between the single smoothing function *v*_*IA*_(*t*) and *LogEnv*(*t*) as defined previously:

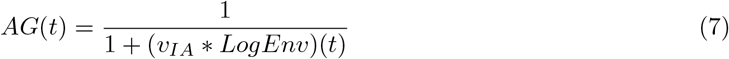

Two essential computational features of the Adaptive Gain function are more evident in this simplified formulation. First, since *v*_*IA*_(*t*) peaks at a time point shortly preceding *t* = 0 and then decays smoothly into the past, the denominator depends on recent sound history (see Supplementary Information for illustrations using values of *τ*_*I*_ and *τ*_*A*_ considered in the Results section). Second, the 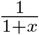nonlinearity constrains Adaptive Gain values to be between 0 and 1, and more importantly, compresses fluctuations in *x* when *x* is large. Thus, when sound levels have recently been high (i.e., (*v*_*IA*_ *∗ LogEnv*)(*t*) is large), Adaptive Gain will be low and small fluctuations in sound level will drive minimal change. However, when sound levels have recently been low (i.e., (*v*_*IA*_ *∗ LogEnv*)(*t*) is small), Adaptive Gain will be high and fluctuations in sound level will have much greater effect on the output. In other words, the 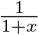nonlinearity imposes a form of intensity gain control. Together, the dependence on recent sound history and the compressive nonlinearity (arising from the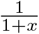 formulation and also from the LogEnv representation) produce dynamic intensity gain control.

Both the dependence on recent sound history and the greater smoothing of fluctuations at high than low sound intensities are evident in comparisons of Adaptive Gain with Envelope and LogEnv representations for example stimuli in Figure 1. To facilitate comparisons between the three representations, Adaptive Gain values were vertically flipped for display (i.e., multiplied by *™*1 and shifted up) and then all three representations were scaled to the same range for visualization, so that the variations of Envelope, LogEnv and Adaptive Gain evolved in the same direction. Note that the Adaptive Gain function not only smooths out sharp changes in sound intensity, illustrating the dependence on recent sound history, but also compresses rapid fluctuations more strongly when sound levels are high than when they are low, enhancing a gain control effect that is also partially achieved by the compressive nonlinearity in the LogEnv transformation alone.

### 2.3 Linear cortical modelling

#### 2.3.1 Forward modelling

The temporal response function *TRF* (*τ*) is the estimated linear response function that transforms a stimulus representation *r*(*t*) into a neural response *n*(*t*) (Lalor et al., 2009), as expressed in the following equation:

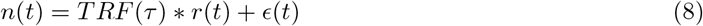

where *∗* is the convolution operator and the error term *ϵ*(*t*) corresponds to the residual neural response that is not explained by the model. Here, *r*(*t*) was either the *Env*(*t*), *LogEnv*(*t*) or *AG*(*t*) representation of the audio signal and TRF modelling was performed separately for each.

Since the relationship between any stimulus feature and the resulting EEG response is not instantaneous, TRFs are always estimated by taking into account multiple time lags. We used a time-lag window for *τ* spanning the interval from -100 ms to 600 ms for all stimulus representations and TRF models.

Estimation of the model *TRF* (*τ*) for each of the three stimulus representations was performed by computing a lagged linear regression between the audio representation *r*(*t*) and the EEG signal *n*(*t*) using the MATLAB mTRF toolbox (Crosse et al., 2016), which incorporates ridge regularization (Wong et al., 2018). Regularization parameters were separately optimised for each model through cross-validation across regularization values in the range [10^*−*10^, 10^10^]. All models were participant-specific, trained and tested on data from only one participant.

### 2.4 Model evaluations

Multiple models were trained for each participant and each audio representation, using segments of 30 seconds of audio along with the corresponding EEG measurements. The training followed a leave-one-out approach, allowing each segment to serve once for testing while the remaining segments were used for training the model. All *n* values (i.e. *n* = 12 for data from Simon et al. (2022) and *n* = 40 for data from Etard and Reichenbach (2022)) of prediction accuracy obtained for one participant were averaged to yield one average prediction accuracy value per participant and audio representation. These values were compared using Wilcoxon signed-rank tests. Paired comparison tests were used to take into account potential variation between participants, due to differences in the amount of data available for training for each dataset (6 minutes available for data from Simon et al. (2024) and 20 minutes for data from Etard et al. (2019)) and also due to individual variability in cortical speech tracking.

The performance of the models was assessed using the Pearson correlation between measured EEG *n*(*t*) and the predicted EEG response *n*^*′*^(*t*) obtained through the convolution of the audio representation *r*(*t*) and the Temporal Response Function *TRF* (*τ*). EEG prediction accuracies will vary across EEG electrode channels depending on how related the data on those channels are to the stimulus representation. The modelling performance on some irrelevant channels may not differ from chance level and might hinder the assessment of models when looking at the prediction accuracy across all channels (Crosse et al., 2021). Here, TRF models computed using different audio representations were compared based on their prediction accuracy for the EEG signal at the FCz channel only. This *a priori* choice of channel was motivated by experimental evidence showing that fronto-central channels exhibit strongest cortical tracking of speech envelope (Drennan and Lalor, 2019; Simon et al., 2024; Zuk et al., 2021; Cantisani et al., 2023).

In addition, TRF weights were used to explore whether spatial and temporal patterns of the predicted EEG signals across electrode positions on the head differed depending on the audio representation. As the ridge regularization factors *λ*, which vary between audio representations and participants, impact the amplitude of the TRF, the TRF weights were z-score normalized for all features and all participants before being averaged across participants for analysis.

#### 2.4.1 Noise floor calculation

To determine the statistical significance of the prediction accuracy and coefficients estimated for the TRF models, we computed noise floors using null models. Null models were created by delaying the audio representation data relative to the EEG data, to ensure that the cortical measurement could not be related to the audio representations when training the null models. The delay was randomly selected from a range between 7.5 and 15 seconds for each leave-one-out iteration of model fitting and cross-validation. All other aspects of the analysis (i.e., use of participant-specific models, cross-validation approach, ridge-regularization parameters, 30 s data segments) were similar for null models and actual models. Results from participant-specific null models were averaged to obtain a noise floor to which prediction accuracy or coefficient values for actual TRF models could be compared.

### 2.5 Data analysis

As some of the data were not normally distributed, non-parametric data analysis was preferred. Wilcoxon rank-sum tests were used to assess if the prediction accuracy distributions differed from the noise floor distributions. Wilcoxon signed-rank tests were used for paired within-subject comparisons between audio representations. All tests were two-tailed, and all results were considered significant with *α* = 0.05. Exact *p*-values are given when higher than 0.0001. Results are shown without Bonferroni correction of *p*-values but all main conclusions were robust to Bonferroni correction.

## 3 Results

### 3.1 Adaptive Gain representation of speech signal improves prediction of cortical speech tracking

We first aimed to determine whether the Adaptive Gain transformation originally derived to improve prediction of mouse auditory thalamic responses to temporally varying sounds could also improve prediction of human EEG responses to continuous speech. For each participant in two independent cortical speech-tracking datasets (Simon et al., 2024; Etard et al., 2019), we estimated temporal response function (TRF) models through linear regression of the EEG signal (at electrode FcZ) with respect to three different transformations of the speech signal: Envelope, LogEnv and Adaptive Gain. TRF models for each of the three audio representations were then evaluated through forward modelling of the EEG signal using segments of continuous speech that had not been used to fit the models (i.e., using cross-validation). Prediction accuracies of the TRF models were quantified with Pearson’s coefficient of correlation between actual and predicted EEG signals.

#### 3.1.1 Pooled data

Analysis of pooled data from both datasets (Figure 2A) showed that prediction accuracies for TRFs estimated using Envelope, LogEnv or Adaptive Gain audio representations were all significantly better than chance (Wilcoxon rank-sum tests, *p <* 0.001 for comparison to noise floor; see Methods for details of noise calculations). However, comparisons between the models revealed that standard Envelope-based TRF models (median prediction accuracy 0.033) were outperformed by both LogEnv models (median = 0.0356; Wilcoxon signed-rank test, *z* = *−*4.1028, *p <* 0.0001, *r* = *−*0.657) and Adaptive Gain models (median = 0.0368; *z* = *−*4.033, *p <* 0.0001, *r* = *−*0.6458). Adaptive Gain models also outperformed LogEnv models (*z* = *−*3.1259, *p* = 0.0018, *r* = *−*0.5005), indicating that logarithmic transformation of the envelope (an initial step in the Adaptive Gain transformation) cannot entirely account for the improvements in prediction accuracy achieved with the Adapative Gain representation.

**Figure 2:**
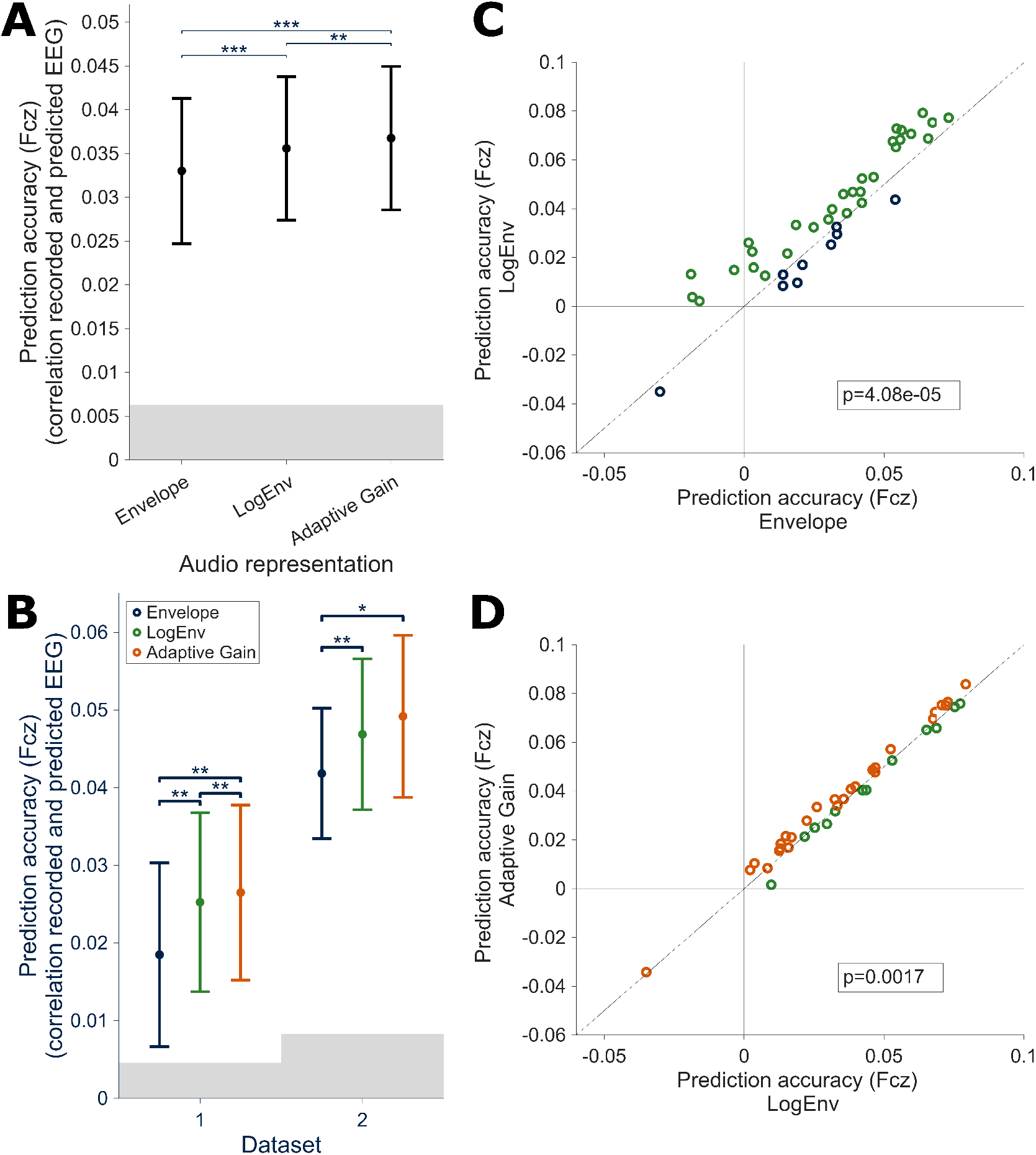
Prediction accuracies for TRF models trained on Envelope, LogEnv, or Adaptive Gain representations of speech. The prediction accuracy corresponds to Pearson’s coefficient of correlation between the measured EEG signal on electrode FcZ and the EEG signal predicted for electrode FcZ based on the incoming speech signal and TRF for each audio representation. **A-B**, Median prediction accuracy. Error bars represent 95% confidence intervals (CI). Gray area corresponds to the noise floor obtained with null models. **A**, Median prediction accuracy for data pooled across the two independent datasets. **B**, Median prediction accuracy for each dataset. Dataset 1 corresponds to Simon et al. (2024) and Dataset 2 corresponds to Etard et al. (2019). **C-D**, Scatterplots comparing prediction accuracy between LogEnv and Envelope representations (C) or between Adaptive Gain and LogEnv representations (D). Each circle corresponds to data from one participant; *p*-values show results of Wilcoxon signed-rank tests. Colours correspond to the legend in B and indicate which of the two audio representations compared in the scatterplot produced higher prediction accuracy for each participant.

These results demonstrate that TRFs estimated using the Adaptive Gain representation of the sound signal predict cortical speech tracking more accurately than those estimated using either the Envelope or the LogEnv representation. Moreover, the results suggest that both components of the Adaptive Gain transformation—logarithmic representation of the stimulus envelope and short-timescale temporal adaptation to sound level—help to improve prediction of cortical speech tracking beyond what is possible with the Envelope representation alone. In control analyses (results not shown), we found that these results were robust to changes in the parameters used for defining the Envelope representation (e.g., 15 Hz instead of 79 Hz low-pass filter).

#### 3.1.2 By dataset

Similar patterns of results were found in each dataset when analyzed separately (Figure 2B). An overall difference in modelling performance was observed between the two datasets included in this study. This difference can be mainly explained by the amount of data available per participant in each dataset.

In Dataset 1 (Simon et al., 2024), TRFs estimated using the Adaptive Gain audio representation (median prediction accuracy = 0.0265) outperformed those estimated using either the Envelope representation (median = 0.0185; Wilcoxon signed rank test, *𝓏* = *−*3.0761, *p* = 0.0021, *r* = *−*0.6712) or the LogEnv representation (median = 0.0253; *𝓏* = *−*2.9718, *p* = 0.0030, *r* = *−*0.6485). The LogEnv representation also yielded significantly better results than the Envelope (*𝓏* = *−*2.9023, *p* = 0.0037, *r* = *−*0.6333).

In Dataset 2 (Etard et al., 2019), TRFs estimated using the Adaptive Gain representation again outperformed those estimated using the Envelope (median prediction accuracy, Adaptive Gain = 0.0492, Envelope = 0.0418; Wilcoxon signed-rank test, *𝓏* = *−*2.3735, *p* = 0.0176, *r* = *−*0.5594). The LogEnv representation (median = 0.0469) also again yielded significantly better TRF results than the Envelope (*𝓏* = *−*2.6783, *p* = 0.0074, *r* = *−*0.6313). For this dataset, the difference in modelling performance between TRFs estimated using Adaptive Gain versus LogEnv audio representations did not reach statistical significance (*𝓏* = *−*1.3718, *p* = 0.1701); however, the trend was consistent with higher prediction accuracy for models based on the Adaptive Gain representation.

#### 3.1.3 By participant

Visualization of data from individual participants in the combined datasets (Figure 2C-D) confirmed that both components of the Adaptive Gain transformation—logarithmic transformation of the envelope and short-timescale temporal adaptation to sound level—improved prediction of cortical speech tracking. Scatterplots comparing prediction accuracies of TRFs estimated using different audio representations showed that modelling performance was improved by the LogEnv relative to Envelope representation for 30 out of 39 participants (Figure 2C, green circles), and improved by the Adaptive Gain relative to the LogEnv representation for 27 out of 39 participants (Figure 2D, orange circles). Notably, the improvements in modelling performance provided by these input transformations were relatively robust to high inter-individual variability in overall TRF modelling success, which is a potentially useful feature given that such variability is common within and across EEG recording studies.

### 3.2 Optimal adaptation time constant for Adaptive Gain representation is longer for humans than mice

The Adaptive Gain transformation was originally derived as part of a temporal processing model optimized for predicting auditory thalamic responses to temporally varying sounds in anesthetized mice (Anderson and Linden, 2016). We wondered if the optimal adaptation time constant *τ*_*A*_ for the Adaptive Gain transformation might be different for predicting cortical speech tracking in awake humans. To address this possibility, we compared the prediction accuracies of TRF models estimated using the Adaptive Gain representation with *τ*_*A*_ values of 5 ms (the minimum, equivalent to the more fundamental integration time constant *τ*_*I*_), 10 ms (the mouse-optimized *τ*_*A*_ value), 25 ms, 50 ms, 75 ms, 100 ms, 250 ms or 500 ms. Prediction accuracies of all models were significantly better than chance (*p <* 0.001), as confirmed by Wilcoxon rank-sum tests comparing model performance to the noise floor. Model performance for the Adaptive Gain representation with the mouse-optimized *τ*_*A*_ value (10 ms) was then compared to model performance for Adaptive Gain representations with all other *τ*_*A*_ values, using Wilcoxon signed-rank tests.

Results showed an increase in model prediction accuracy as the value of *τ*_*A*_ increased until it reached a maximum for *τ*_*A*_ = 100 ms, then decreased for longer values of *τ*_*A*_ (see Figure 3A). Prediction accuracy was significantly higher than that achieved with the mouse-optimized *τ*_*A*_ = 10 ms (median = 0.0368) for *τ*_*A*_ = 25 ms (median = 0.0381; Wilcoxon signed-rank test, *𝓏* = *−*4.7726, *p <*.0001, *r* = *−*0.7642), *τ*_*A*_ = 50 ms (median = 0.0402; *𝓏* = *−*4.6331, *p <*.0001, *r* = *−*0.7419), *τ*_*A*_ = 75 ms (median = 0.0419; *𝓏* = *−*4.3679, *p <*.0001, *r* = 0.6994), and *τ*_*A*_ = 100 ms (median = 0.0431; *𝓏* = *−*3.7539, *p <*.000141, *r* = 0.6011). However, prediction accuracy for *τ*_*A*_ = 5 ms (median = 0.0365) was lower than for the original mouse-optimized parameter (*𝓏* = 4.3679, *p <*.0001, *r* = *−*0.6994). Model prediction accuracy was not significantly different from that obtained with the original mouse-optimized *τ*_*A*_ = 10 ms for *τ*_*A*_ = 250 ms (median = 0.0418; *𝓏* = *−*1.9397, *p* = 0.0524) and *τ*_*A*_ = 500 ms (median = 0.0406; *𝓏* = *−*0.8233, *p* = 0.4103).

**Figure 3:**
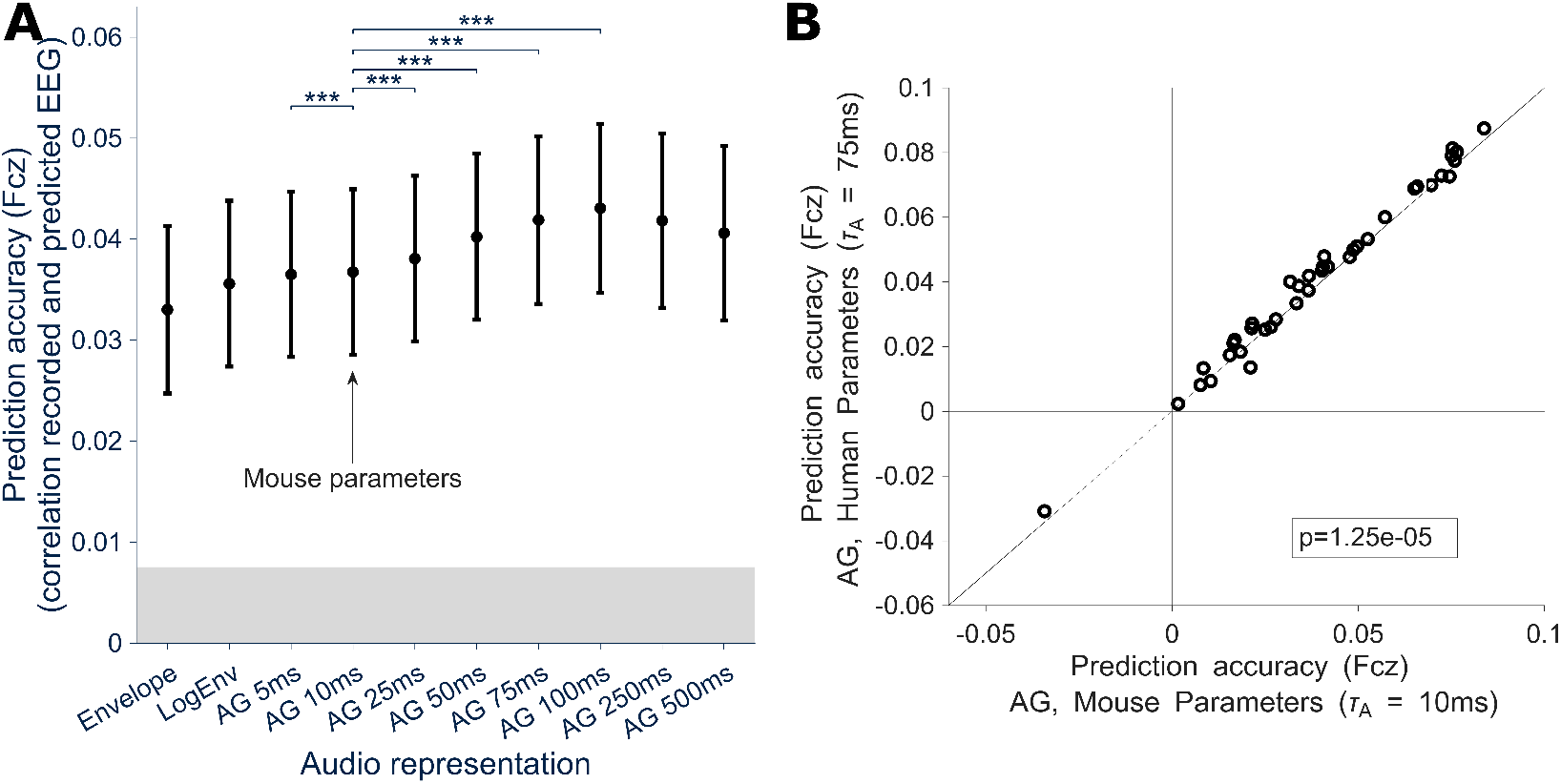
Prediction accuracies for TRF models trained on Envelope, LogEnv and Adaptive Gain representations of speech with varying *τ*_*A*_ values. Conventions as in Figure 2. **A**, Median prediction accuracy *±*95% CI. Brackets and asterisks indicate significant differences in prediction accuracy between TRF models trained on Adaptive Gain representations with the mouse-optimized value for the adaptation time constant *τ*_*A*_ versus other values (Wilcoxon signed-rank tests, *\* * *p <* 0.001). **B**, Scatterplot comparing TRF prediction accuracy for the Adaptive Gain representation with a human-optimized adaptation time constant (*τ*_*A*_ = 75 ms) versus mouse-optimized (*τ*_*A*_ = 10 ms) parameters. Each circle corresponds to data from one participant; *p*-value shows result of Wilcoxon signed-rank test.

These results indicate that the Adaptive Gain transformation is more useful for predicting cortical speech tracking in humans with an adaptation time constant *τ*_*A*_ of 25-100 ms rather than the *τ*_*A*_ = 10 ms value originally derived in studies of anesthetized mice. We chose *τ*_*A*_ = 75 ms as a human-optimized value for further investigations, since time constants of 50, 75 and 100 ms all produced models with prediction accuracy statistically indistinguishable from the peak. As illustrated by the fact that almost all data points are above the diagonal line in Figure 3B, model prediction accuracy was indeed higher for nearly all participants when using this human-optimized *τ*_*A*_ value rather than the original mouse-optimized value.

### 3.3 Topographical maps and TRF shapes suggest that Envelope, LogEnv and Adaptive Gain representations capture similar speech-tracking mechanisms

The Adaptive Gain transformation was developed to capture biologically relevant adaptive normalization in time. Our hypothesis is that the Adaptive Gain transformation suppresses and enhances features in the speech signal that would also be suppressed or enhanced by central auditory temporal processing mechanisms. This hypothesis assumes that the underlying speech-tracking mechanisms modelled with TRFs using the Envelope, LogEnv, and Adaptive Gain representations as regressors should be qualitatively similar, even if model performance is quantitatively different. To test this hypothesis, we compared topographic maps of prediction accuracy and the TRF shapes (i.e., the weights of the TRF models) obtained using the different audio representations as regressors.

Figure 4 compares the maps of the prediction accuracy obtained with models trained on Envelope and LogEnv representations and Adaptive Gain representations either with the original adaptation time constant from mouse studies (*τ*_*A*_ = 10 ms) or the human-optimized adaptation time constant identified here (*τ*_*A*_ = 75 ms). All four maps present a similar pattern of increased prediction accuracy on the frontocentral electrodes, with lateralization on the right hemisphere and improved prediction accuracy also on the left occipital electrodes. This pattern is consistent with previous work on auditory TRFs modelled on the stimulus envelope or representations of slow temporal amplitude variations in sound (Simon et al., 2024; Hjortkjær et al., 2020; Prinsloo and Lalor, 2022; Di Liberto et al., 2020; Lindboom et al., 2023; Borges et al., 2025; Drennan and Lalor, 2019). The similarity of patterns here suggests that the mechanisms modelled with the four audio representations are comparable. On the electrodes that are best predicted with all models, an increase of prediction accuracy can be seen and follows the trend observed on FCz, emphasising that using the Adaptive Gain representation, especially with a longer length of the adaptive window, leads to a better-performing model than using the Envelope or the LogEnv representations.

**Figure 4:**
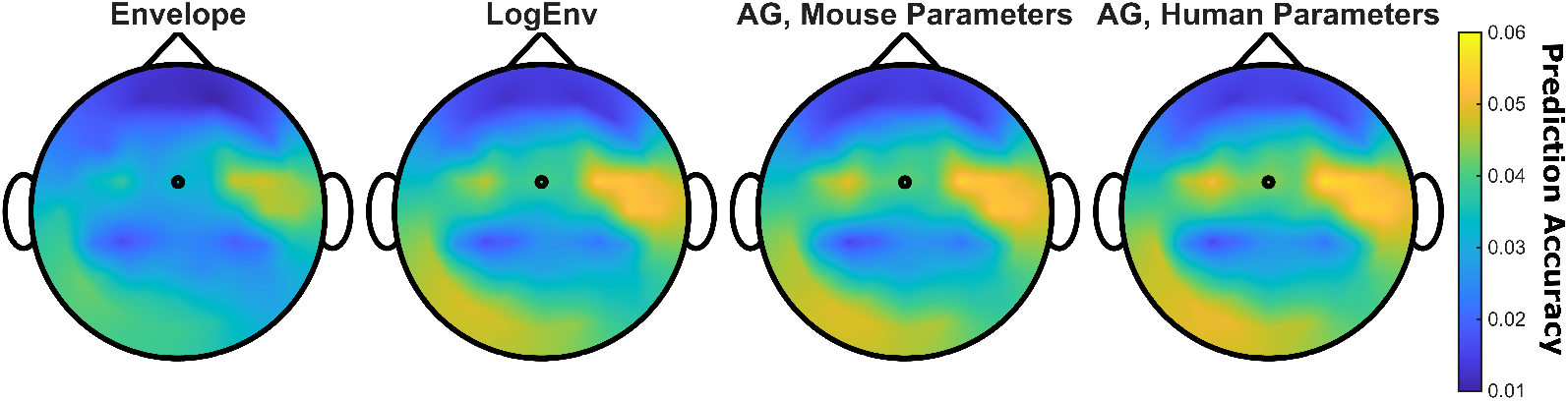
Topographical maps of TRF model prediction accuracy for each audio representation and for each EEG electrode. Black circles correspond to the position of electrode FcZ.

The weights of the models, which represent the shape of the Temporal Response Functions (TRFs), also show notable similarities (Figure 5). For all four audio representations analyzed, the TRFs recorded at the FCz electrode exhibit a typical P50-N100-P200 complex, a pattern associated with auditory event-related potentials. The topographic mapping of the TRF models indicates that this activity is primarily concentrated around the fronto-central electrodes, which has also been observed in other studies (Simon et al., 2024; Hjortkjær et al., 2020; Di Liberto et al., 2020; Lindboom et al., 2023; Dieudonné et al., 2025; Carta et al., 2023; Borges et al., 2025). However, there are differences in both amplitude and latency. The peaks are more defined, with a higher amplitude for the TRFs derived from the Adaptive Gain representations compared to the Envelope representation. Analysis of the latency of the peaks reveals a consistent time shift for the two Adaptive Gain conditions. This time shift is likely attributable to the Adaptive Gain transformation itself, and specifically the delay due to smoothing with the function *v*_*IA*_ (see Supplementary Information for details).

**Figure 5:**
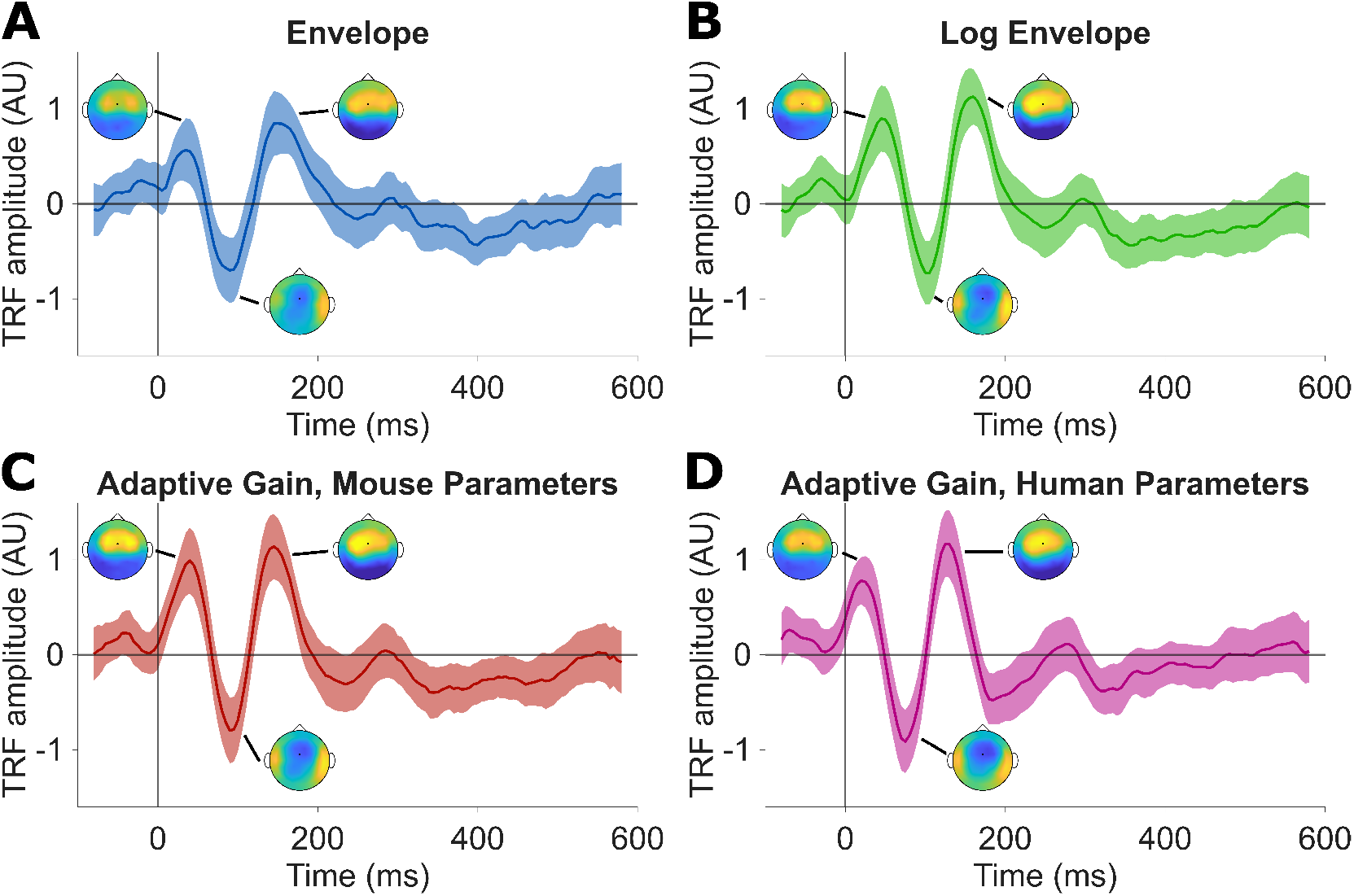
Temporal response functions (TRFs) derived from linear regression between EEG data and the different representations of the speech stimuli: Envelope (A), LogEnv (B), Adaptive Gain with mouse-optimized parameters (C) and Adaptive Gain with human-optimized parameters (D). Shaded areas correspond to 95% CI. Topographic maps show the spatial distribution of the activities for the P50, N100 and P200 components.

### 3.4 Adaptive Gain representation also improves prediction of cortical speech tracking for not-understood speech

Assuming the Adaptive Gain transformation captures fundamental mechanisms of temporal processing in the central auditory system shared by mice and humans (with possibly different time constants), then it should to improve cortical speech tracking for speech that was not understood as well as for understood speech. We tested this hypothesis using remaining data from both datasets, in which participants listened to speech in languages they did not understand (Figure 6). Results were very similar to those obtained for cortical speech tracking of understood languages (compare to Figure 2A and Figure 3A). Again, time constants of 50, 75 and 100 ms all produced models with prediction accuracy statistically indistinguishable from the peak (although here the peak median prediction accuracy occurred at *τ*_*A*_ = 50 ms instead of *τ*_*A*_ = 100 ms in Figure 3A). We did not pursue a more detailed comparison of results for understood and not-understood languages because of the difficulty of controlling for levels of attentional engagement. However, we note that TRF prediction accuracies were very similar for understood and not-understood languages.

**Figure 6:**
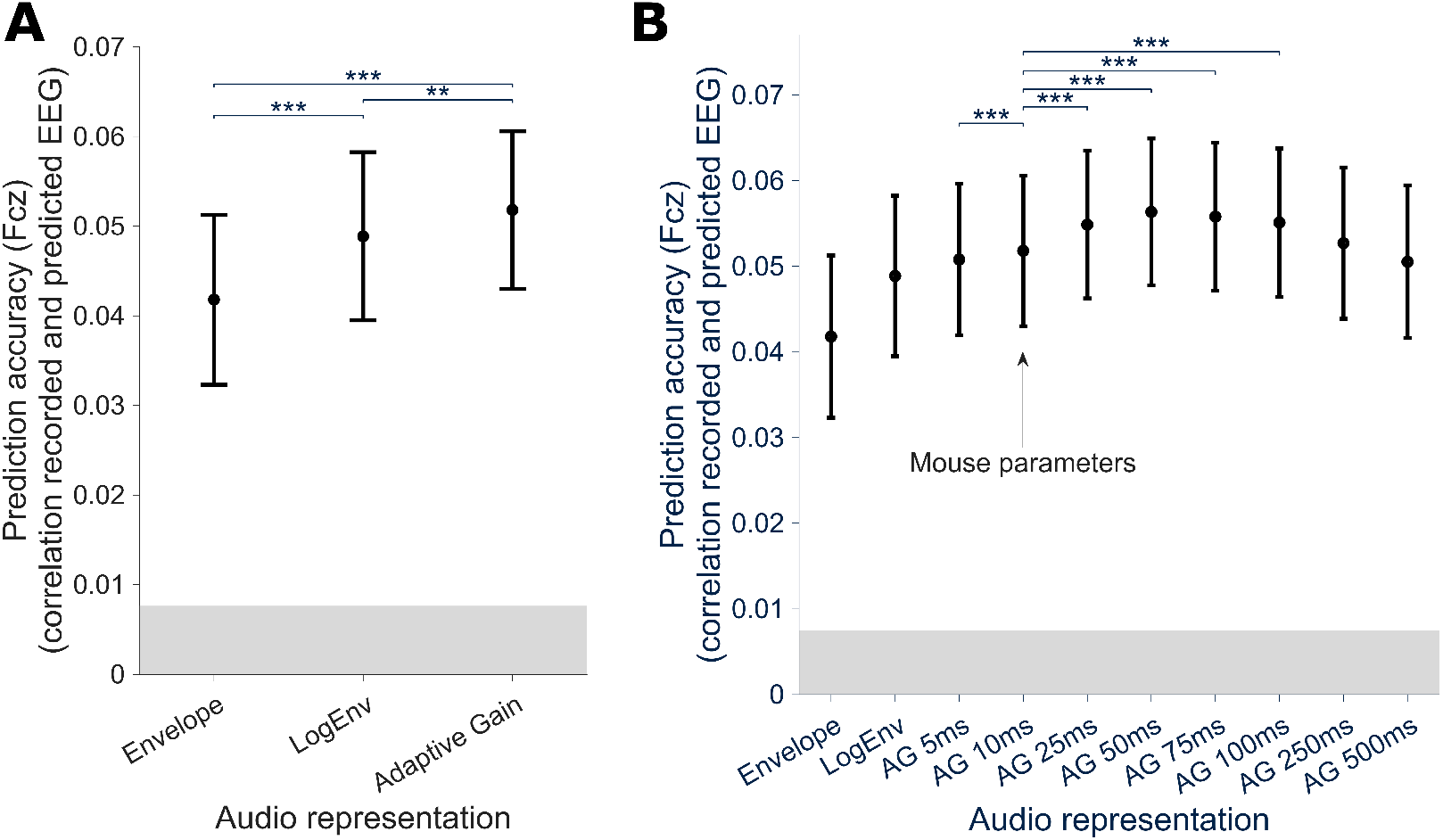
Prediction accuracies obtained on the remaining data of both datasets, in which participants listened to speech in not-understood languages (Finnish for data from Simon et al. (2022) and Dutch for data from Etard and Reichenbach (2022)). Conventions as in Figure 2 and Figure 3. **A**, Median prediction accuracy *±*95% CI for TRF models trained on Envelope, LogEnv and Adaptive Gain representations with the original mouse-optimized parameters (compare with Figure 2A). **B**, Median prediction accuracy *±* 95% CI for Envelope and LogEnv representations and for Adaptive Gain representations with varying *τ*_*A*_ values (compare with Figure 3A).

This result demonstrates that the Adaptive Gain transformation improves prediction of cortical speech tracking for not-understood as well as understood speech.

## 4 Discussion

This study investigated whether a simple Adaptive Gain mechanism previously identified in studies of central auditory temporal processing in mice could be used to improve prediction of human cortical responses to continuous speech. The results demonstrate that Temporal Response Function (TRF) models estimated using the Adaptive Gain representation of speech outperform models estimated using either the standard Envelope or the logarithmic envelope (LogEnv) representations, indicating that adaptive mechanisms play a significant role in shaping cortical tracking of speech. Moreover, parameter optimization for human EEG revealed that longer adaptation time constants (around 50–100 ms, compared to 10 ms in Anderson and Linden (2016)) yielded the best prediction performance, suggesting that the temporal dynamics of adaptation in the auditory cortex of awake humans differ from those previously observed in the auditory thalamus of anesthetized mice.

These findings extend previous work showing that the use of biologically grounded auditory transformations improves the performance of cortical speech-tracking models (Lindboom et al., 2023; Biesmans et al., 2016; Drennan and Lalor, 2019). Here, the biologically grounded transformations were nonlinearities in intensity and temporal processing: logarithmic representation of sound intensity (which compressed and smoothed amplitude variations in the sound envelope) and short-timescale adaptive normalization in time (which enhanced responsiveness to sound features occurring when sound levels had recently been low, and suppressed responsiveness to sound features occurring when sound levels had recently been high).

The improvements in TRF modelling performance observed with the Adaptive Gain over the Envelope representation arose from both nonlinearities, as was illustrated by the intermediate improvements achieved with TRF models based on the LogEnv representation alone. These findings are biologically plausible and consistent with previous results from animal and human studies. There is extensive psychophysical and neurophysiological evidence for logarithmic coding of sound intensity in the auditory cortex (e.g. Aiken and Picton (2008); Ahrens et al. (2008)), although previous work on linear modelling of cortical speech tracking has not found that adding logarithmic scaling to the envelope improves performance (Biesmans et al., 2016, 2015; Drennan and Lalor, 2019). The adaptive component of the Adaptive Gain transformation is a novelty for cortical speech tracking studies but also biologically plausible, as it approximates short-timescale neural adaptation processes that are ubiquitous in the auditory pathway (Anderson and Linden, 2016; Kopp-Scheinpflug and Linden, 2021). Previous studies in animal models have found that similar adaptive mechanisms improve the prediction of single-unit responses to temporally varying sounds in the auditory cortex (Willmore et al., 2016). Here, we observed improvements in prediction of cortical speech tracking with the Adaptive Gain transformation compared to both Envelope and LogEnv representations across two independent datasets (Etard and Reichenbach, 2022; Simon et al., 2022), highlighting the robustness of the findings.

Analysis of topographical maps of prediction accuracy and TRF shapes also provided evidence that the Adaptive Gain representation is an improvement over the Envelope representation. The topographic distribution of TRF prediction accuracy for the Envelope representation, with maximal prediction accuracy observed on fronto-central electrodes, was consistent with results reported in previous envelope-tracking studies (Simon et al., 2024; Prinsloo and Lalor, 2022; Hjortkjær et al., 2020; Borges et al., 2025; Drennan and Lalor, 2019). Similar topographical distributions of TRF prediction accuracy were observed for models based on the LogEnv and Adaptive Gain representations, suggesting similarity in the underlying speech-tracking mechanism detected by the models. The similar shape of the TRFs obtained using the Envelope, LogEnv and Adaptive Gain representations also supports this interpretation. For all audio representations, a three-peaked complex, resembling the typical P50-N100-P200 complex found in auditory event-related potentials, can be clearly observed in the TRF morphology, and was also concentrated on the fronto-central electrodes. The slight latency shift observed in the TRFs estimated using the Adaptive Gain representations, relative to the TRFs estimated using the Envelope and LogEnv representations, is consistent with the temporal smoothing and delay introduced by the double convolution with integration and adaptation time windows in the Adap-tive Gain computation (see Supplementary Information). Taken together, these results support the idea that the Adaptive Gain transformation of speech enhances, rather than replaces, Envelope-based models of slow cortical tracking.

Adaptive mechanisms have been found throughout the auditory pathway, operating over a range of different time scales (Robinson and McAlpine, 2009; Pérez-González and Malmierca, 2014; King and Walker, 2020; Shamma and Fritz, 2014; Willmore and King, 2023). Animal studies have revealed that while some adaptive mechanisms are driven by subcortical processing, including at the level of the thalamus (Anderson et al., 2009), additional adaptive mechanisms arise in the auditory cortex on a longer time scale (Asari and Zador, 2009; Ulanovsky et al., 2004; Dean et al., 2008). These observations suggest that the longer adaptation time constant for the human-optimized than mouse-optimized Adaptive Gain transformation might reflect differences between auditory cortex and thalamus (and between speech processing and processing of simpler temporally varying sounds) as well as differences between awake humans and anesthetized mice. Further studies could explore how dynamic adaptation in different frequency channels might relate to cortical speech tracking, by applying the Adaptive Gain transformation separately to sound signals in different frequency bands (instead of only to the broadband sound waveform as done here). Another possible extension would be to integrate the Adaptive Gain audio representation with linear modelling of subcortical as well as cortical responses during speech listening (Kulasingham et al., 2024; Etard et al., 2019), to examine the contributions of subcortical versus cortical adaptation to speech tracking in humans.

In conclusion: the Adaptive Gain representation is a simple and easily implemented alternative to the Envelope representation that reliably improves TRF modelling performance for continuous speech, demonstrating a role for adaptive temporal normalization in cortical speech tracking. Notably, the Adaptive Gain representation improved prediction of cortical speech tracking even with mouse-optimized parameters, and for not-understood as well as understood speech, suggesting that it captures fundamental properties of central auditory temporal processing. Therefore, the Adaptive Gain representation might be useful as an alternative to the Envelope representation to improve predictions of human EEG responses not only for speech but for any temporally varying sounds. This improvement could translate into practical benefits, for example in applications such as auditory attention decoding in multi-speaker environments or development of novel objective measures for auditory processing disorders.

## Acknowledgements

This work was funded by a research grant to J.F.L. and M.C. from the National Institute for Health and Care Research / University College London Hospitals Biomedical Research Centre (NIHR-UCLH BRC) “Hearing Health” Theme.

## Supplementary Information

### 1 Form of the Adaptive Gain term

The Adaptive Gain model applies two successive convolutions to the logarithm of the stimulus envelope (LogEnv) and then applies a nonlinear transformation 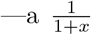 inversion intended to compress and constrain output values between 0 and 1. Here we attempt to clarify and simplify the Adaptive Gain function by reducing the two successive convolutions to a single smoothing function.

The Adaptive Gain function as defined in Anderson and Linden (2016) is as follows.

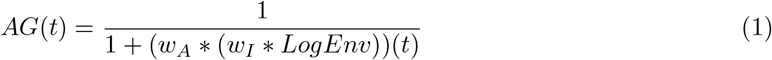

where *\** signifies convolution, *w*_*A*_ and *w*_*I*_ are the exponential weight functions detailed in the main text of the current paper, and LogEnv is the logarithm of the stimulus envelope as also defined in the main text.

Convolution (discrete and continuous) is both commutative:

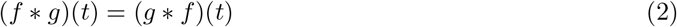

and associative:

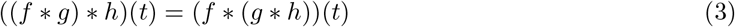

So using (3) and (2) in (1):

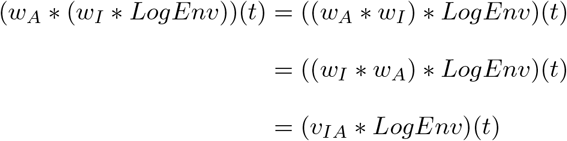

To give a closed form for *v*_*IA*_ we perform a continuous convolution, before resampling back to discrete functions. This gives the same result as discrete convolution provided sampling is fast enough (double the highest relevant frequency in the sound pressure waveform *s*(*t*), which here would be the upper frequency limit for human hearing). The limits here are given under the assumption that the weight functions *w*_*I*_ and *w*_*A*_ continue to *∞*. Error terms from truncation and discretization will be discussed and calculated subsequently.

Substituting in continuous formulations of the definitions of *w*_*I*_ and *w*_*A*_ given in Anderson and Linden (2016) with appropriate normalizing constants 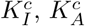 for the continuous case:

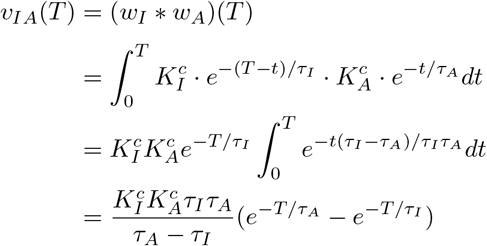

We absorb the constant term in the above formulation into a normalization constant 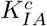. It may seem as if we’ve introduced an asymmetry with the negative sign, but in fact as *e*^*−T/τ A*^ *≥ e*^*−T/τ I*^ *⇔ τ*_*A*_ *≥ τ*_*I*_, we have the sign in the numerator always cancelling the sign in the denominator of 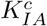, maintaining the symmetry we expect from (2). Thus without loss of generality we assume *τ*_*I*_ *≤ τ*_*A*_ and 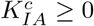 to give the closed form:

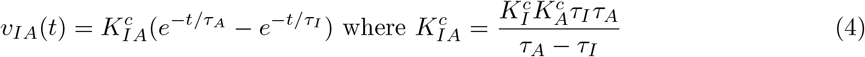

In the continuous case, the constants are:

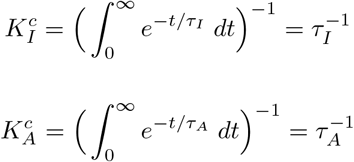

Therefore 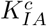 simplifies to 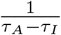, and we obtain simplified forms for *v*_*IA*_(*t*) and the Adaptive Gain function *AG*(*t*):

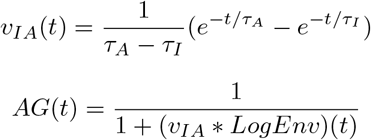

### 2 Compression effect of the non-linear 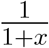 transformation

The 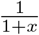 transformation compresses variation in *v*_*IA*_ *\*LogEnv*(*t*) at higher amplitudes (as the logarithmic transformation also does for the stimulus envelope). The 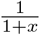 transformation of *v*_*IA*_ *\* LogEnv*(*t*) also ensures that *AG*(*t*) is bounded in the interval (0, 1] (for positive input, which *v*_*IA*_ *\* LogEnv*(*t*) always is).

The practical consequence of this 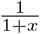 nonlinearity (in combination with peak delay in *v*_*IA*_) is that when sound levels have recently been very high, further increases in volume will have minimal effect on Adaptive Gain, as it will already be near zero producing maximal damping. Conversely, when sound levels have recently been very low, sudden increases in volume will have much more impact as Adaptive Gain will be close to one.

### 3 Error terms from discretization and truncation of *v*_*IA*_

#### 3.1 Discretization

As stated above (see “Form of the Adaptive Gain term”), provided sampling is fast enough for our use case (human hearing) we have no relevant discretization error between performing the convolution discretely, or continuously and then resampling. The sampling performed is at 48 kHz with a period of *δ ≈* 0.02 ms, which is fast enough to achieve this (the accepted sample rate for human audio being 44.1 kHz).

Thus the only discretization error present is error between the continuous integral of *v*_*IA*_ and the discrete integral of *v*_*IA*_. This error would only be relevant if the continuously calculated constant 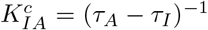 were used. In practice, *v*_*IA*_ is only used discretely and is normalized to make the discrete integral 1, avoiding discretization error.

#### 3.2 Truncation of *w*_*I*_, *w*_*A*_ **and** *v*_*IA*_

In the main text, we use *w*_*A*_ and *w*_*I*_ functions that are cut off to 0 at *t*-values bigger than 5*τ*_*A*_ and 5*τ*_*I*_. The rationale for this cutoff is that it is 4*σ* away from the mean of the exponential decay functions *w*_*I*_ and *w*_*A*_, and thus the error from this cutoff is *<* 0.01.

To achieve the same small error, we borrow some ideas from probability theory, as the function *v*_*IA*_ is the probability density function of the sum of two independent exponential random variables. So the mean of *v*_*IA*_ is *µ*_1_ + *µ*_2_, where *µ*_1_, *µ*_2_ are the means of the two exponential random variables. Now as we are taking the sum of two exponential random variables, we would expect the variation of this sum to be smaller than the variation of a single exponential random variable with the same mean as the sum given. So qualitatively, we expect the cutoff point sufficient to reduce the error to *<* 0.01 in the exponential case to reduce the error to *<* 0.01 in the sum-of-exponentials case. The sufficient cutoff for the exponential case is 5*µ* where *µ* is the mean of the random variable. So using this cutoff for *v*_*IA*_ – 5(*τ*_*I*_ + *τ*_*A*_) – should be sufficent to produce less error.

Indeed we can calculate the error in all cases used in the paper and see it is always smaller than 0.01, shown below via plot of error against *x* = *τ*_*A*_*/τ*_*I*_:

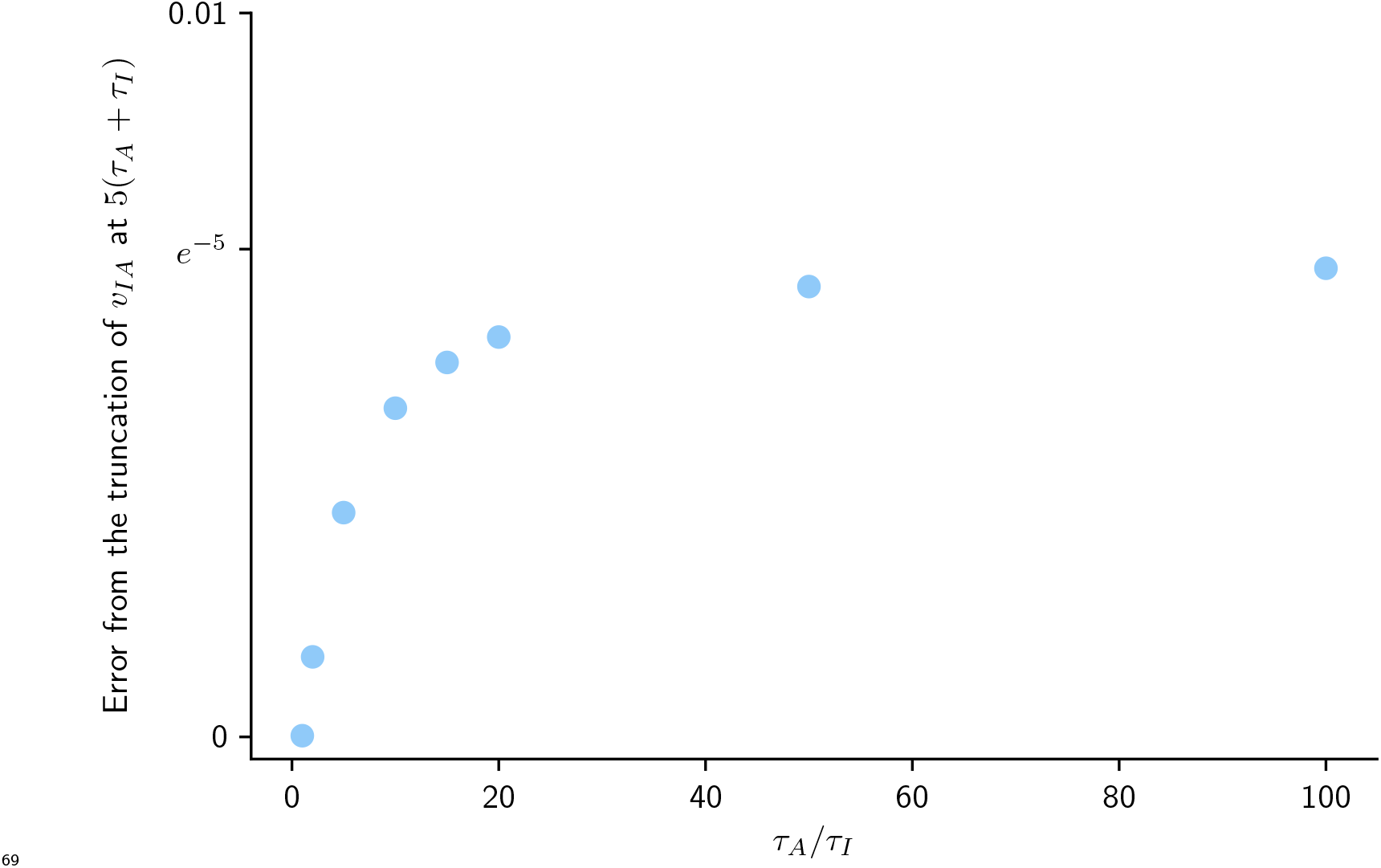

### 4 Character of *v*_*IA*_

We compare the shape of *v*_*IA*_ for different values of *τ*_*A*_ given a fixed *τ*_*I*_ = 5 ms:

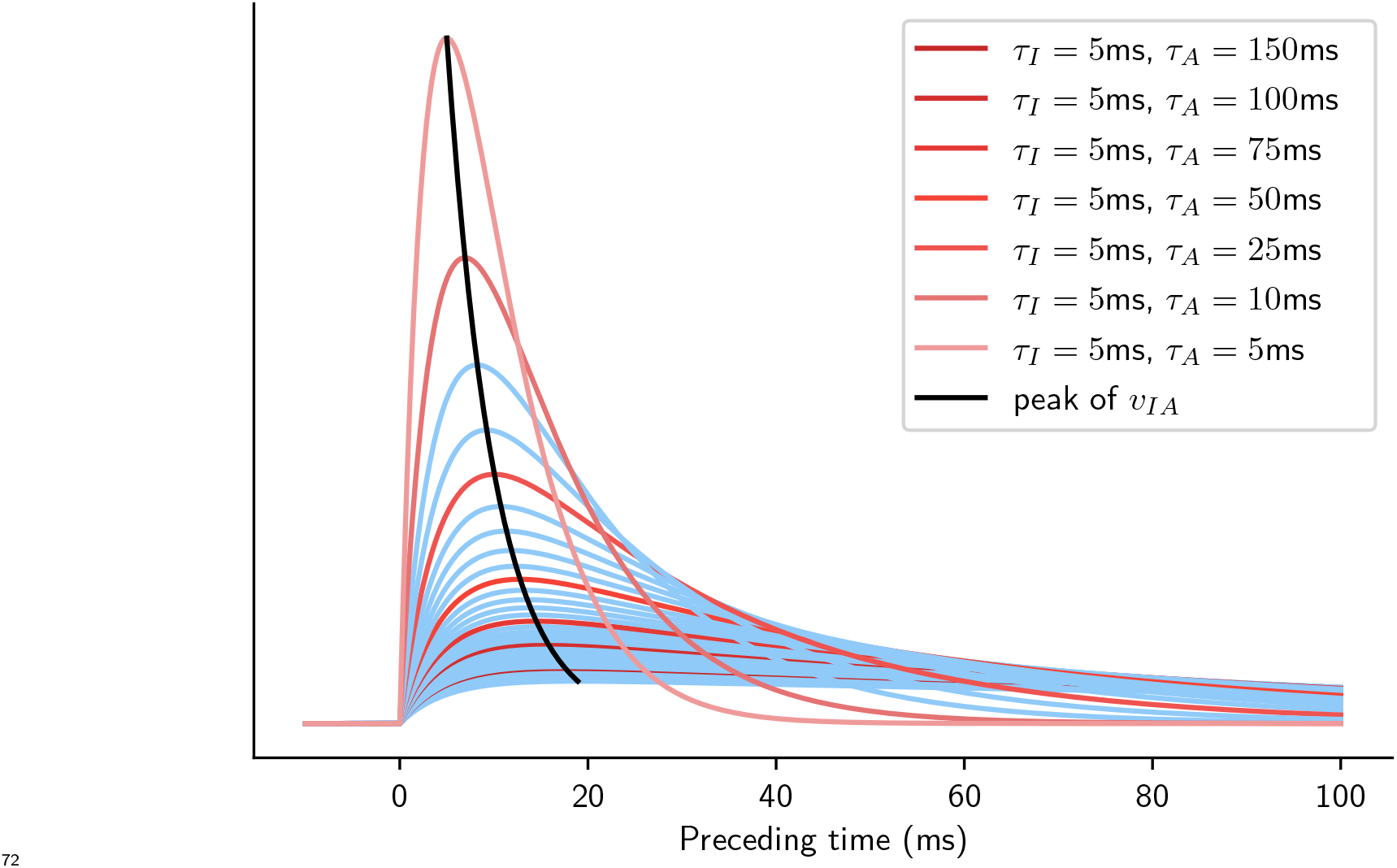

The major difference between a standard exponential smoother and the *v*_*IA*_ smoothing function used here is the delay in the peak. In a standard smoother, the peak is at *t* = 0, whereas here we can see the peak occurs here at *t* = 10 *−* 20 ms. When this function is convolved with the logarithm of the stimulus envelope *LogEnv*(*t*) in the adaptive gain calculation, this peak delay means that the maximal contribution to the final convolution at a point *T* comes from the point in *LogEnv*(*t*) with *t ≈ T* = 10–20 ms for *τ*_*I*_ = 5 ms and *τ*_*A*_ values as indicated in the legend above. This is in contrast to the standard smoother which gives maximal contribution from the input at the same point as where the convolution is evaluated. This delay is biologically plausible given that the shortest auditory cortical response latencies typically fall in a similar range.

